# A dual-fluorescent recombinant for live observation of Herpes simplex-type 1 infection outcomes

**DOI:** 10.1101/2023.06.07.544108

**Authors:** Luke F. Domanico, G. P. Dunn, O. Kobiler, M.P. Taylor

**Author notes:** Corresponding Author: Matthew P. Taylor, Ph.D. Office.

## Abstract

Critical stages of lytic Herpes simplex type 1 (HSV-1) replication are marked by the sequential expression of immediate early (IE) to early (E), then late (L) viral genes. HSV-1 also persists in neuronal tissues via a non-replicative, transcriptionally repressed infection called latency. Understanding the regulation of lytic and latent transcriptional profiles provides focused insight into HSV-1 infection and disease. We sought a fluorescence-based approach to observe temporal progression of HSV-1 infection at the single-cell level. We constructed and characterized a novel HSV-1 recombinant that reports IE and L gene expression by fluorescent protein detection. The dual-reporter HSV-1 visualizes IE gene expression by a CMV promotor-driven YFP, and L gene expression by an endogenous mCherry-VP26 fusion. We confirmed that viral gene expression, replication and spread of infection in epithelial cells is not altered by the incorporation of the fluorescent reporters. Interference with viral DNA polymerase activity abolishes VP26-mCherry detection late in HSV-1 infection, visually reporting the failure to complete viral replication. Viral replication in primary neurons is not altered, but retrograde-directed inoculation of the dual-reporter HSV-1 exhibits a modest reduction in titer, compared to unlabeled HSV-1. Low-dose axonal inoculation in the presence of small molecule modulation of neuronal signaling results in divergent outcomes of YFP and mCherry detection, suggesting different states of latent and lytic replication. Rigorous characterization of this dual-reporter HSV-1 recombinant has demonstrated the utility of temporal observation of HSV-1 replication in live cells and the potential for further insight into the dynamics of infection.

**Importance:** Herpes simplex virus-type 1 (HSV-1) is a prevalent human pathogen that infects approximately 67% of the global population. HSV-1 invades the peripheral nervous system, where latent HSV-1 infection persists within the host for life. Immunological evasion, viral persistence, and herpetic pathologies are determined by regulation of HSV-1 gene expression. Studying HSV-1 gene expression during neuronal infection is challenging but essential for the development of antiviral therapeutics and interventions. We constructed and characterized a dual-fluorescent HSV-1 recombinant that enables visualization of IE and L gene expression. The recombinant HSV-1 is used to observe the progression and outcome of infection. We demonstrate that drug treatments targeting cellular pathways can manipulate latent HSV-1 infection in neurons to achieve divergent outcomes of infection.

## Introduction

Viral replication is a regulated process determined by the coordination of temporally defined events. The regulation of HSV-1 gene expression plays an important role in the progression of viral replication and the outcome of infection. During lytic HSV-1 replication, viral genes are expressed in a regulated cascade of three classes of genes: immediate-early (IE), early (E) and late (L) (1). The IE genes are expressed first from the viral genome, yielding multifunctional proteins that establish a permissive cellular environment supportive of HSV-1 replication (2). E gene expression is promoted by IE proteins and includes the viral DNA replication machinery. Replication of the HSV-1 genome stimulates L gene expression, which include many structural proteins and correlates with assembly of progeny virions (2). Initial lytic HSV-1 replication occurs in epithelial tissues, before spreading to innervating sensory neurons (3). In neurons, HSV-1 establishes a persistent, transcriptionally suppressed infection alternative to lytic replication, termed latency. The latently infected neuron provides a lifelong viral reservoir that can reactivate, causing subclinical herpetic lesions or fatal Herpes Simplex encephalitis. Investigation of HSV-1 gene expression in epithelial and neuronal cells has contributed much to our understanding of HSV-1 disease and therapeutic intervention (4-9). However, the cellular and viral factors involved in the establishment of latency remain elusive.

To investigate viral gene expression, replication, and spread, many researchers employ HSV-1 recombinants that express various reporter proteins (8, 9). Fluorescent protein-expressing reporter viruses have been employed to visually observe HSV-1 infection (10-21). HSV-1 isolates have been engineered with fluorescent protein fusions to viral proteins of interest (20), or expression cassettes under regulation of a chosen viral promotor (22) or inserted within a targeted genomic locus (10, 14, 15). Importantly, fluorescent reporter viruses are employed to study lytic gene expression (16), assembly and spread (14,23, 24) and latent HSV-1 infection (17, 18 22, 25). Only recently have fluorescent reporters been used to investigate the time-dependent intricacies of infection at the single cell level (16, 26).

Recent advances in molecular, biochemical and imaging methods have enabled single-cell analyses that provide high-resolution observation of cellular and viral factors influencing HSV-1 infection (13, 27, 28), as well as outcomes of infection that are underrepresented in population-level analyses (19, 27, 29). However, these approaches use methods that are not amenable to sequential observation of individual cells through the course of HSV-1 replication. Many single-cell HSV-1 studies capture endpoints of viral replication, yielding limited interpretations on the progression of infection. As a result, the dynamic progression of HSV-1 infection is missed. Therefore, tools and methods enabling temporal observation of HSV-1 gene expression provide critical insight into the progression of HSV-1 replication.

In this study, we evaluate an HSV-1 recombinant that encodes a CMV-driven eYFP to report IE gene expression, and an mCherry fusion to endogenous VP26 to report L gene expression. This dual reporter virus enables temporal visualization of HSV-1 replication. The work here compares replicative competence and gene expression of the recombinant HSV-1 to the unlabeled, parental strain. We demonstrate that the dual-fluorescent reporter virus retains replication and gene expression kinetics comparable to wild-type HSV-1 infection in epithelial cells and primary neurons. We then demonstrate that differential detection of fluorescent reporters correlates with inhibition of viral replication by antiviral treatment. We employed a model of silent neuronal infection to identify altered transcriptional profiles induced by small molecule treatments. This critical characterization demonstrates the utility of the dual-fluorescent reporter virus to monitor viral replication progression and provide significant insight into the events of HSV-1 gene expression and replication.

## Results

### Comparison of dual reporter recombinant HSV-1 replication

We have isolated a recombinant HSV-1 that possesses two fluorescent reporter proteins, enabling real-time visualization of viral gene expression and replication (**fig. 1A**). To report IE gene expression, a yellow fluorescent protein (eYFP) with a membrane-targeting motif under the regulation of a CMV promotor is inserted between UL37 and UL38 open reading frames of the HSV-1 genome (14). To report L gene expression, an in-frame red fluorescent protein (mCherry) fusion to the N-terminus of the minor capsid protein VP26 (UL35) was generated in the endogenous UL35 locus (30, 31). Vero cells infected with the dual reporter HSV-1 produce diffuse cytosolic YFP with enrichment at the cell periphery, and nuclear-localized puncta of mCherry (**fig. 1B**). To assess the suitability of the recombinant HSV-1 as a reporter of HSV-1 replication, we compared viral replication kinetics of unlabeled, parental HSV-1 strain17 (unlabeled HSV-1), and dual reporter HSV-1. Control infections with single-reporter recombinants possessing only the CMV-driven eYFP (HSV-1 YFP) or the mCherry-VP26 fusion (HSV-1 mCherry-VP26), were also evaluated by single-step replication. Similar extents of infectious virus are produced at all timepoints evaluated. This result demonstrates that the presence of either fluorescent reporter alone, or in combination, does not hinder replication of dual reporter HSV-1 (**fig. 1C**). We next compared intercellular spread of dual reporter HSV-1 to unlabeled HSV-1 by quantifying plaque size. At 72 hours post infection (hpi) monolayers were imaged for endogenous fluorescence (**fig. 1D**), then fixed and stained with polyclonal HSV anti-serum to compare plaque diameter (**fig. 1E**). The dual reporter HSV-1 produced larger average plaque area (4.7x10^5^ μm^2^) compared to unlabeled HSV-1 (3.9x10^5^ μm^2^) but with greater plaque size diversity (**fig. 1F**). Together, these results demonstrate that the dual reporter HSV-1 exhibits no deleterious defect on viral replication or intercellular spread in epithelial cells.

**Figure 1:**
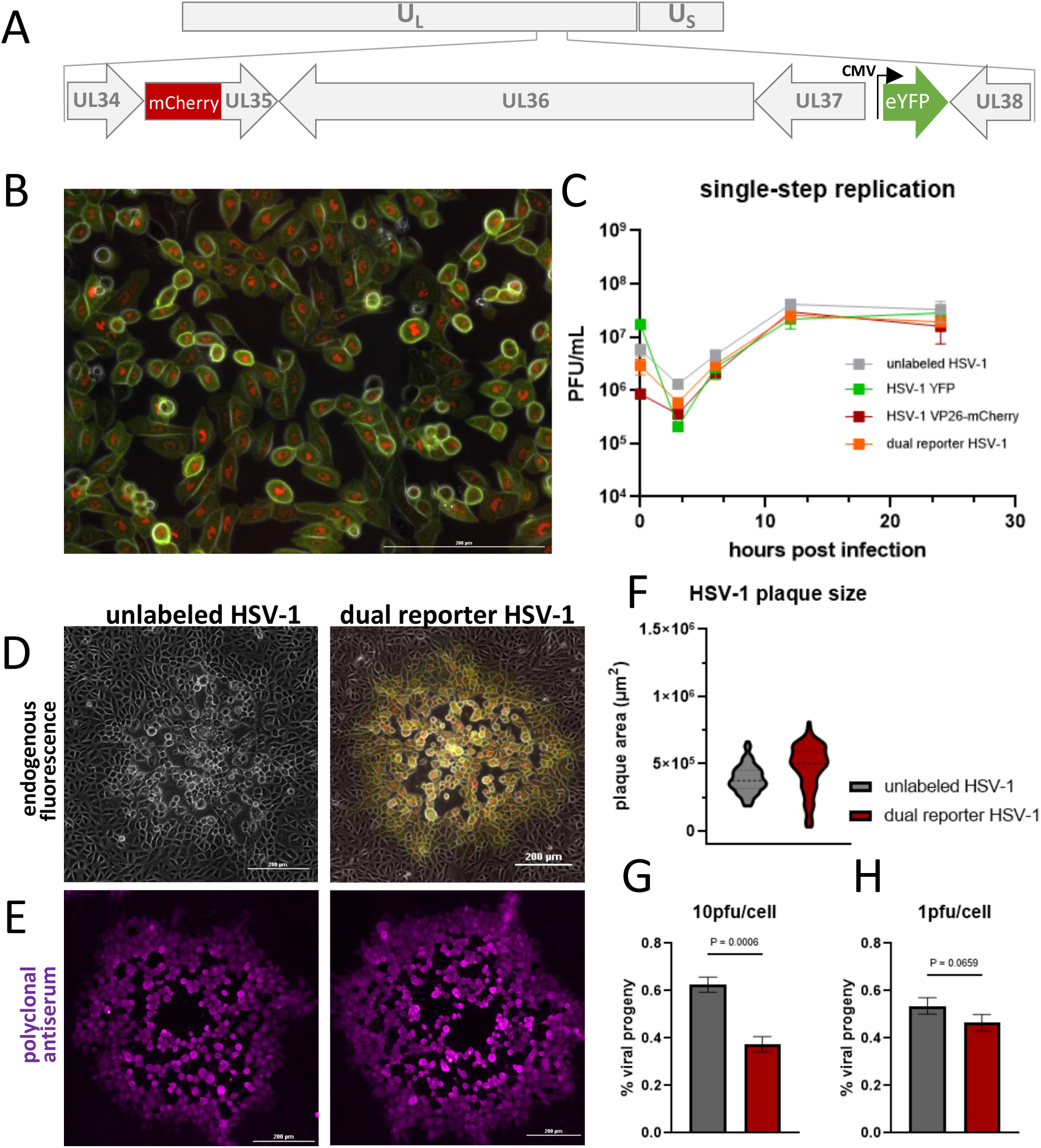
The dual reporter HSV-1 demonstrates wild-type replication and fitness. **A.)** Schematic of the dual reporter HSV-1 genome. **B.)** Image of Vero cells infected with OK15 at 10pfu/cell, imaged 8hpi. **C.)** Growth curve of unlabeled HSV-1, HSV-1 YFP, HSV-1 VP26-mCherry and dual reporter HSV-1. Vero cells inoculated with virus at 10pfu/cell, harvested at 0, 3, 6, 12, and 24hpi for plaque assay. Data representative triplicate infection, each titered in duplicate. Error represented as SD. **D.)** Representative images of unlabeled HSV-1 or dual reporter HSV-1 plaques, imaged at 72hpi. Images are a 3-channel merge of YFP (green) and mCherry (red) fluorescence overlayed on a phase contrast image (grayscale) **E.)** Unlabeled HSV-1 or dual reporter HSV-1 plaques were fixed and stained with polyclonal HSV serum (Dako). Immuno-staining is false-colored purple. **(F)**. Quantification of plaque area from immuno-stained monolayers infected with unlabeled HSV-1 or dual reporter HSV-1. Measurements made using NIS Elements software, data representative n=200 plaques per virus, infections performed in duplicate. **G.)** Vero cells were coinfected at 10 pfu/cell simultaneously with unlabeled HSV-1 and dual labeled HSV-1, harvested at 8hpi and plaque assayed for progeny quantification **(H.)** Vero cells were coinfected at 10 pfu/cell simultaneously with unlabeled HSV-1 and dual labeled HSV-1, harvested at 8hpi and plaque assayed for progeny quantification. Data representative of two independent coinfections, each assayed in duplicate. Error represented as SD.

HSV-1 with CMV-driven fluorescent cassettes have been reported to exhibit moderately hindered viral replicative capacity, as measured by a co-infection competition assay (21). To test the capacity for cellular competition between unlabeled HSV-1 and the dual reporter HSV-1, cells were infected simultaneously with both viruses. The replicative capacity of each virus was assessed by quantifying viral progeny by plaque fluorescence. The dual reporter HSV-1 demonstrated a modest reduction in progeny compared to unlabeled HSV-1, with fluorescent plaques constituting approximately 40% of progeny following high multiplicity of infection (MOI) (**fig. 1G**). When inoculated under limiting MOI conditions, we observed a reduced but insignificant difference in the number of dual reporter HSV-1 plaques compared to unlabeled HSV-1 progeny (**fig. 1H**). From this evaluation, we can affirm that the dual reporter HSV-1 possesses a comparable replicative capacity to other recombinant viruses and unlabeled HSV-1 (21).

### Dual reporter HSV-1 fluorescence and viral gene expression over time

Our overall goal was to construct and isolate a recombinant HSV-1 reporter virus that enables observation of HSV-1 replication progression. To visualize the progression of viral replication, we infected Vero cells at high MOI to observe (**fig. 2A-C**) the timing of and quantify (**fig. 2D**) fluorescent protein detection via live-cell fluorescence microscopy. At 1 hour post infection (1hpi), we do not detect YFP or mCherry signal (**fig. 2A and 2D**). By 4hpi, YFP signal was detectable, while mCherry signal was still not detectable (**fig. 2B and 2D**). At 6hpi, YFP detection has increased, and onset of mCherry signal is observed in some cells (**2D and supplemental movie 1**). There is variability in the onset of detectable mCherry expression across the population of infected cells. By 8hpi, we detected robust YFP and mCherry signal, visually reporting the achievement of fulminant HSV-1 infection (**fig. 2C and 2D**).

**Figure 2:**
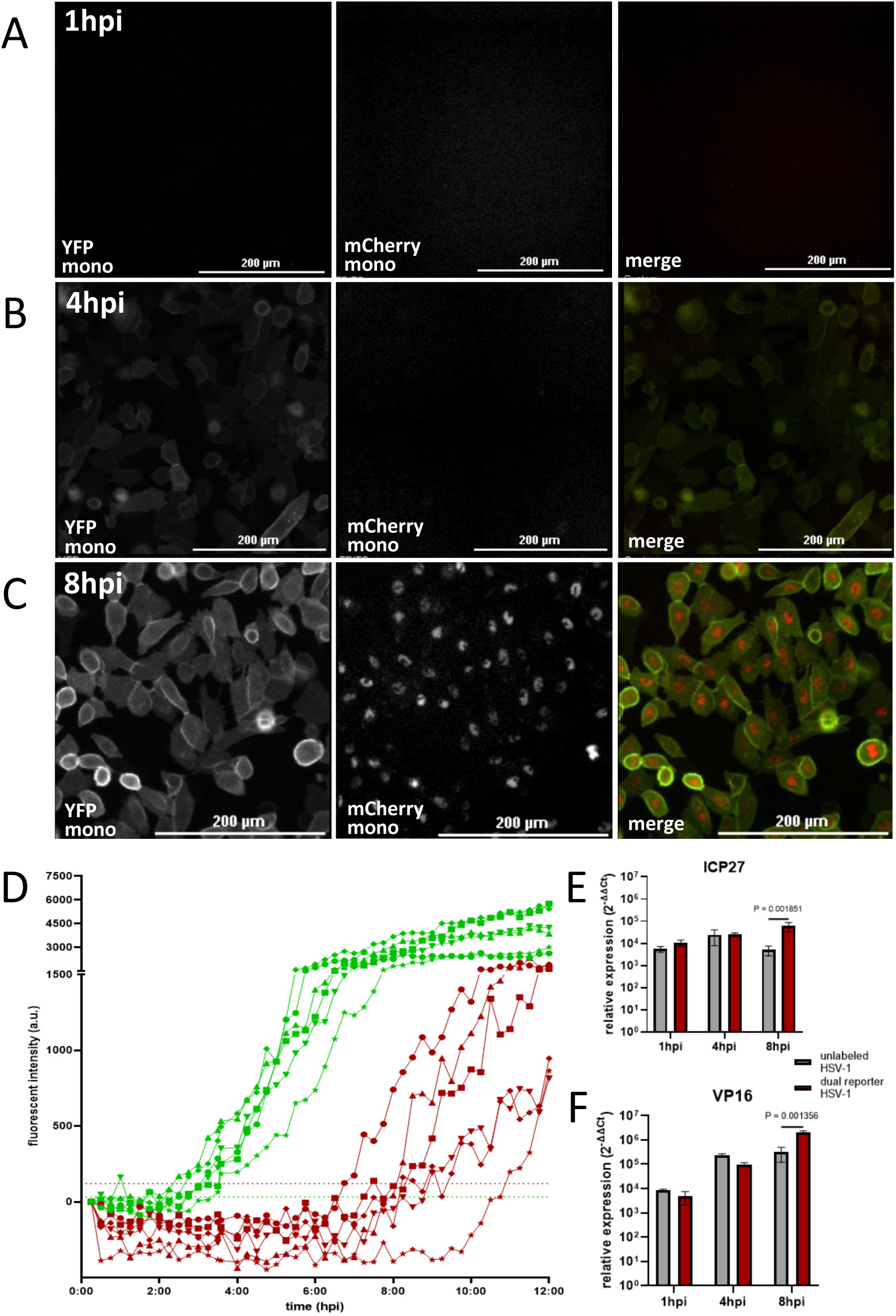
The dual reporter HSV-1 demonstrates representative HSV-1 viral gene expression kinetics. **A.)** Images of Vero cells infected with dual reporter HSV-1 at 10pfu/cell, imaged at 1, 4 and 8hpi. All images acquired with equal exposure time and power and represented with equal display parameters. YFP and mCherry fluorescence are presented in monochrome for each channel and in color (green and red, respectively) for the merged image. **B.)** Plot of fluorescent intensity measurements from cells infected with dual reporter HSV-1 over time. Quantification performed using Fiji ImageJ. Threshold values set at 32 a.u. for YFP detection, and 120 a.u. for mCherry detection. Presented are fluorescent intensity traces for YFP (green) and mCherry (red) for 6 individual cells, labeled as different shapes. **C and D.)** Vero cells infected with unlabeled HSV-1 (grey) or dual reporter HSV-1 (red) at 10pfu/cell and harvested for RT-qPCR at 1, 4 and 8 hpi. Relative quantitation of RNA performed using ΔΔCt method. 28s rRNA used as endogenous control. qPCR data representative of three technical replicates. Comparison performed via multiple unpaired t-tests; error represented as SD.

It is critical that the dual reporter HSV-1 does not exhibit delayed gene expression. We sought to temporally assess viral gene expression by comparing viral transcription between unlabeled HSV-1 and the dual reporter HSV-1. Vero cells were infected and harvested at sequential timepoints post infection for RT-qPCR of viral transcripts. First, we chose HSV-1 ICP27 as a target to evaluate IE gene expression. ICP27 is a multifunctional protein encoded by the UL54 gene that is expressed with IE kinetics and is maintained throughout infection. ICP27 is critical regulator of cellular and viral gene expression (32, 33). Overall, we detect ICP27 transcripts at 1hpi, increasing at 4hpi that is then maintained at 8hpi (**fig. 2E**). Interestingly, we observed increased detection of ICP27 in dual reporter HSV-1 infected cells compared to unlabeled HSV-1 infected cells at 8hpi (**fig. 2E**).

Next, we sought to assess dual reporter HSV-1 expression of an L transcript. VP16 is a structural protein encoded by the UL48 gene that is incorporated in the tegument layer of HSV-1 virions. VP16 is critical for the regulation of IE genes and is expressed with true-late kinetics (34). We observed increased detection of VP16 from 1hpi through 8hpi in cells infected with dual reporter HSV-1 or unlabeled HSV-1 (**fig. 2F**). From 1hpi to 8hpi, VP16 mRNA increased 100-fold in cells infected with either virus. However, similar to ICP27 detection, we detected significantly more VP16 at 8hpi in dual reporter HSV-1-infected cells compared to cells infected with unlabeled HSV-1 (**fig. 2F**). The differences in viral transcript detection, particularly at 8hpi are difficult to account for, yet do not detract from the comparability of viral transcription at the earlier, critical timepoints assessed by the fluorescent reporters. This data demonstrates that HSV-1-expressed fluorescent reporter detection temporally reports HSV-1 IE and L gene expression without a deleterious effect on endogenous viral transcription.

### Inhibition of viral DNA replication impacts fluorescent protein detection

HSV-1 L gene expression is stimulated by duplication of the viral genome (1,2). Many studies have evaluated the inhibition of HSV-1 DNA duplication to thwart viral replication (23). We reasoned that inhibiting genome replication in infected cells would yield YFP detection, but diminished or absent mCherry detection at 8hpi. To test the hypothesized presentation of fluorescence, cells were inoculated with dual reporter HSV-1 then treated with inhibitors of viral DNA replication one hour later. We used the antiviral drug acyclovir (Acv) to interfere with viral DNA elongation (35), or phosphonoacetic acid (PAA) to inhibit viral DNA polymerase activity (36). At 8hpi, infected cells treated with DMSO control or either antiviral exhibited a similar level of YFP detection (**fig. 3A**). However, infected cells treated with Acv (**fig. 3B**) or PAA (**fig. 3C**) demonstrate a marked decrease in mCherry detection. PAA treatment had a greater impact on reducing mCherry detection compared to Acv treatment, likely due to more effective interference of HSV-1 genome duplication.

**Figure 3:**
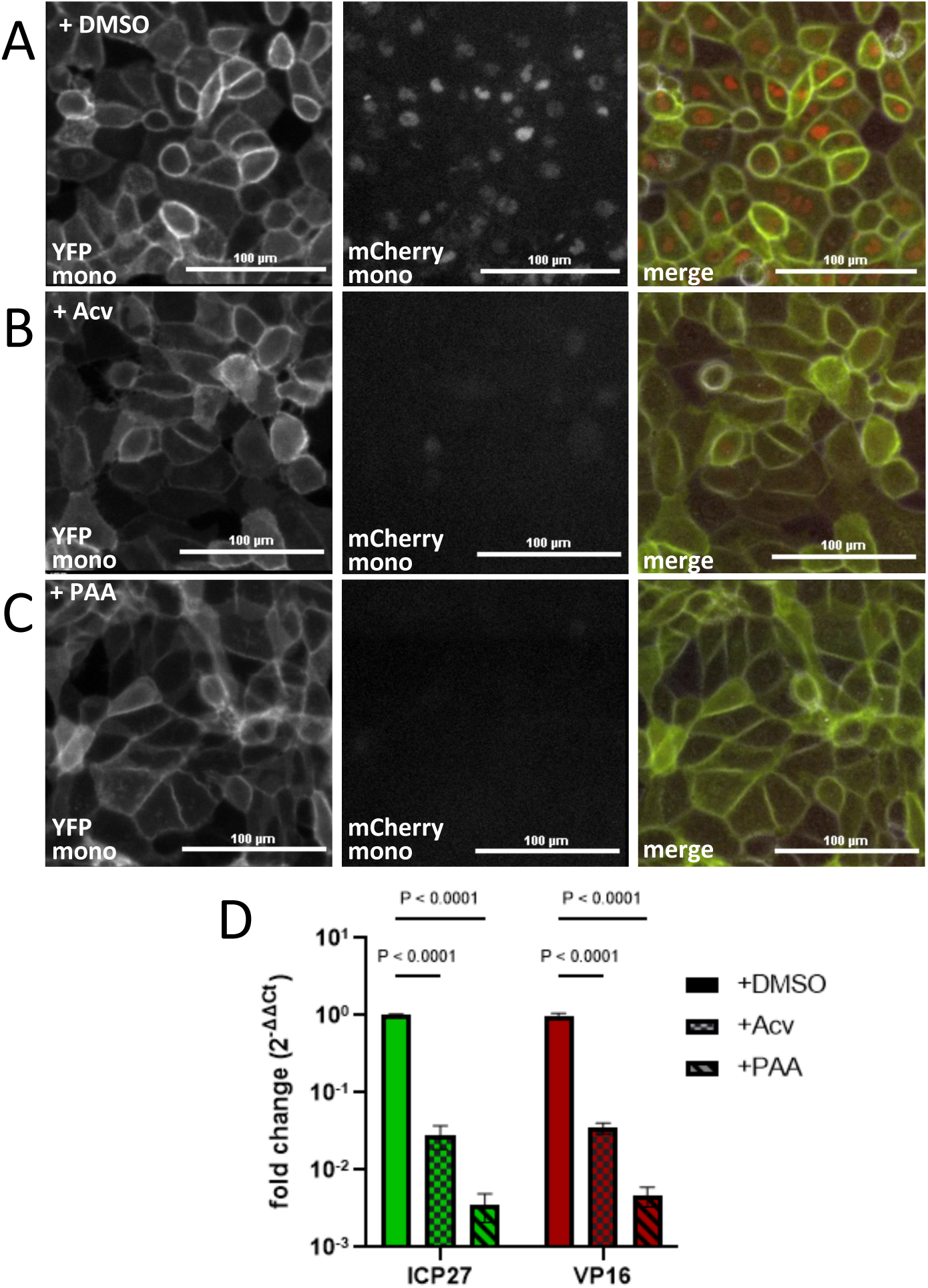
Inhibition of dual reporter HSV-1 infection demonstrates representative FP detection. Vero cells infected with dual reporter HSV-1 at 10 pfu/cell and imaged at 8hpi **A.)** in the presence of 10uM DMSO **B).** in the presence of 10uM acyclovir **C).** or 400ng/mL phosphonoacetic acid. All images acquired with equal exposure time and power and represented with equal display parameters. YFP and mCherry fluorescence are presented in monochrome for each channel and in color (green and red, respectively) for the merged image. **D.)** Parallel infections were performed and harvested for RT-qPCR. Relative quantitation of RNA performed using ΔΔCt method. 28s rRNA used as endogenous control. qPCR data representative of three technical replicates. Comparisons performed via two-way ANOVA using Tukey’s multiple comparison test, error represented as SD.

To correlate the effect of antiviral treatment on fluorescent protein detection and endogenous viral gene expression, parallel infections were analyzed by RT-qPCR for viral transcript detection. Analysis of dual reporter HSV-1 gene expression at 8hpi demonstrated marked reduction in ICP27 and VP16 in the presence of Acv or PAA, compared to DMSO control (**fig. 3D**). PAA treatment elicited a greater reduction in ICP27 and VP16 expression than Acv, consistent with the detection of the mCherry reporter. These results demonstrate that HSV-1 expressed YFP and mCherry detection reliably reports the extent of viral gene expression in the presence of antiviral inhibitors.

### Evaluation of dual reporter HSV-1 neuronal infection

Investigating HSV-1 infection of the nervous system is critical to understanding and preventing viral disease. To evaluate the capacity of dual reporter HSV-1 to infect, traffic and replicate in neurons, we employed a modified compartmentalized primary mouse superior cervical ganglia (SCG) neuron culturing model (37, 38). This model spatially organizes neurons in a physiologically representative manner such that infection can be targeted to neuronal axons or soma by inoculation of the respective compartments (**fig. 4A**). Importantly, similar models have been employed to analyze the capacity for HSV-1 to undergo retrograde infection and establish viral latency (22, 39). Compartmentalized SCG neurons inoculated with dual reporter HSV-1 produce somatic YFP detection that extends along axons and nuclear-localized mCherry puncta (**fig. 4A**).

**Figure 4:**
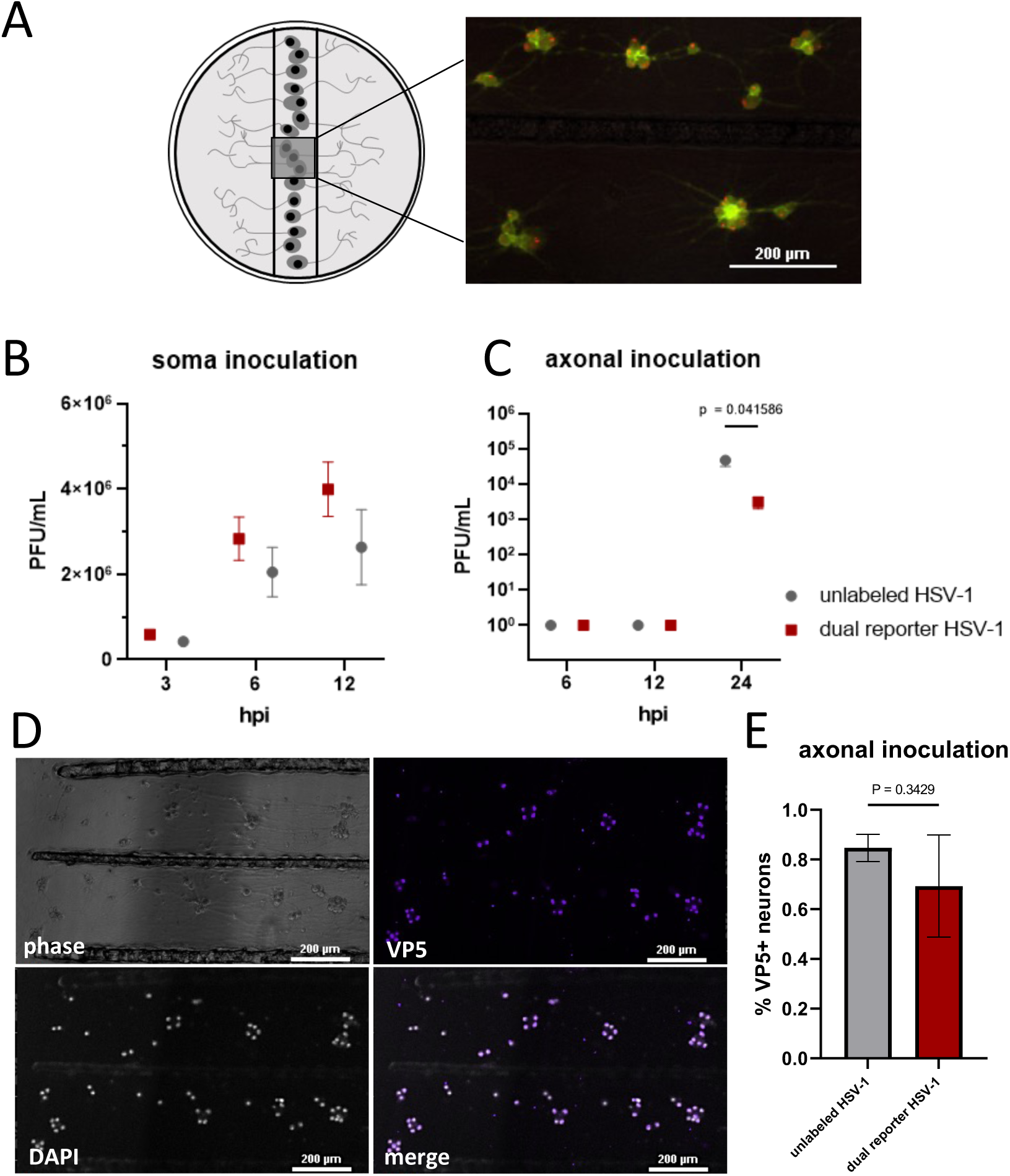
OK15 demonstrates representative HSV-1 neurotropism of compartmentalized SCGs. **A.)** Schematic of compartmentalized neuron culturing system for experiments presented in panels B-E. **B.)** Image of compartmentalized neuronal soma infected with dual reporter HSV-1 upon axonal inoculation, imaged at 24hpi. Presented is a two-channel merge of YFP (green) and mCherry (red) fluorescence **C.)** Compartmentalized SCG neurons were inoculated with unlabeled or dual reporter HSV-1 on neuronal soma and were harvested at indicated timepoints for plaque assay. Data representative of three neuronal infection replicates, each titered in duplicate. Error represented as SD. **D.)** Compartmentalized SCG neurons were inoculated with unlabeled or dual reporter HSV-1 on neuronal axons and were harvested at indicated timepoints for plaque assay. Data representative of three neuronal infection replicates, each titered in duplicate. Error represented as SD. **E.)** Compartmentalized SCG neurons were inoculated on neuronal axons and were fixed and stained for HSV-1 VP5. Representative images of compartmentalized SCG neurons inoculated with dual reporter HSV-1 on neuronal axons that were fixed and stained at 24hpi. Presented is the Phase contrast image (grey), VP5 monoclonal antibody (purple), nuclear counterstain (white), and a two-channel merge image of the fluorescence. **F.)** Quantification of VP5+ SCG neurons performed using NIS elements. Neuronal nuclei identified and counted via Hoechst nuclear counterstain, and VP5 positivity was subsequently quantified. Immunofluorescence data representative of four neuronal infection replicates, error represented as SD. All microscopy images acquired with equal exposure time, power and represented with equal channel intensity parameters.

To examine neuronal replication of the dual reporter HSV-1, single-step replication of HSV-1 infection of neurons was performed. Here, neuronal soma in the middle-compartment were inoculated with unlabeled HSV-1 or dual reporter HSV-1 and harvested at indicated timepoints for plaque assay to assess viral titer. Across all sampled timepoints, the dual reporter HSV-1 exhibited a statistically insignificant enhancement in neuronal replication compared to unlabeled HSV-1 (**fig. 4B**).

Next, we sought to assess neuronal infection from retrograde-directed inoculation of the dual reporter HSV-1. Axons in the lateral compartments were inoculated with unlabeled HSV-1 or dual reporter HSV-1, and subsequently neuronal soma were harvested for plaque assay to quantify viral titer. At 6hpi and 12hpi, no viral titer was recovered from SCG neurons infected with either unlabeled or dual reporter HSV-1 (**fig. 4C**). At 24hpi, when an increase in titer was detected, dual-reporter HSV-1 titer was slightly reduced (9-fold) compared to unlabeled HSV-1 **(fig. 4C**). To understand if the observed reduction in titer is explained by a difference in the number of infected neurons, we evaluated the frequency of neuronal infection. Following the same axonal inoculation, neurons were fixed at 24hpi and immuno-stained for detection of HSV-1 VP5 (**fig. 4D**). VP5 stained soma were quantified and represented as percent of total neurons in each culture. We observed a statistically insignificant decrease in the percentage of VP5-positive neurons following dual reporter HSV-1 infection compared to unlabeled HSV-1 (**fig. 4E**). These experiments suggest that dual reporter HSV-1 is capable of neuronal infection and replication but has a modest reduction in neuroinvasive capacity compared to unlabeled HSV-1, potentially at the step of retrograde-directed trafficking.

### Manipulating silent infection results in divergent outcomes of gene expression

The cellular and environmental factors determining the establishment of latent HSV-1 infection are not understood. Multiple reports demonstrate that a low-dose axonal inoculation promotes the establishment of viral latency without the use of antivirals (22, 39). We employed our dual reporter HSV-1 to assess viral gene expression during the establishment of viral latency. To achieve a latent HSV-1 infection, we applied a limiting inoculating dose to axons in the lateral compartments of our culturing system and imaged neuronal soma every 24 hours. Through 120hpi, we observed no YFP or mCherry signal in neuronal cultures, suggesting a silent neuronal infection of the dual-reporter HSV-1 (**fig. 5A**).

**Figure 5.**
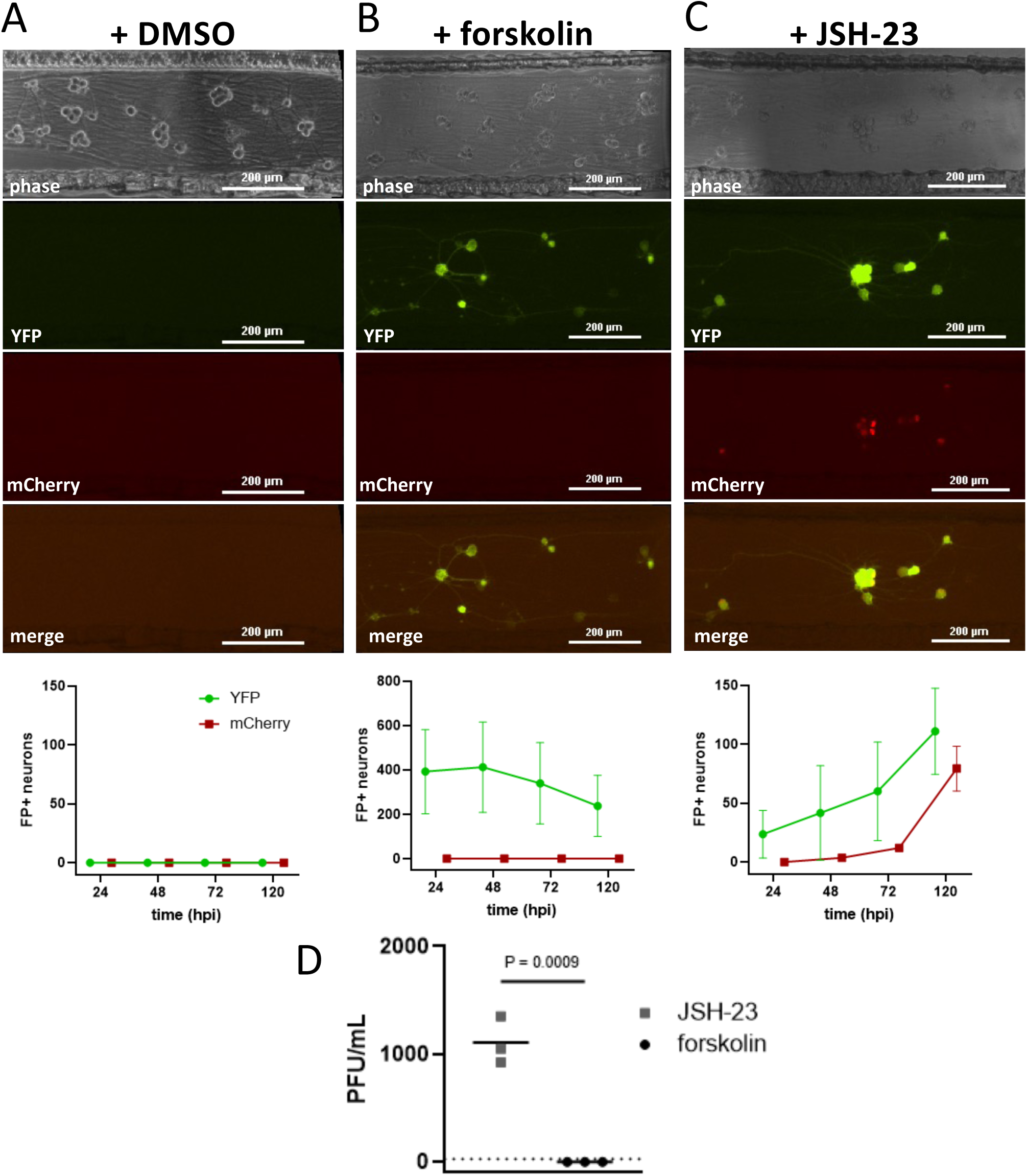
Low-dose axonal inoculation reports divergent outcomes of HSV-1 infection. **A.)** Images of compartmentalized neuronal soma infected with dual reporter HSV-1 via low-dose axonal inoculation in the presence of **A.)** DMSO, **B.)** forskolin or **C.)** JSH-23. Neuron cell bodies were imaged at 72hpi. Presented are representative images with phase contrast (grey), YFP (green), mCherry (red), and two-channel fluorescence merge for each condition. Cell counts of YFP+ and mCherry+ soma for the 4 days after inoculation are below each image. All microscopy images acquired with equal exposure time, power and represented with equal channel intensity parameters. Data representative of three neuronal infection replicates per treatment condition, error represented as SD. **D.)** Compartmentalized neuronal cultures from panels **B. and C.** were harvested at 120hpi for plaque assay. Horizontal bar represents mean, comparison performed via one-way ANOVA.

We hypothesized that manipulating neuronal signaling could alter the establishment of a silent infection, resulting in YFP and mCherry detection. To manipulate the outcome of our low-dose axonal inoculation, neuronal media was supplemented with either JSH-23 or forskolin at the time of axonal inoculation. JSH-23 is an inhibitor of NF-kB translocation that diminishes nuclear accumulation of p65 in response to HSV-1 infection (40). Forskolin stimulates the production of adenylyl cyclase and has conventionally been used to reactivate latent HSV-1 infection *in vitro* (41-44). Neurons supplemented with JSH-23 at the time of low-dose axonal inoculation produced detectable YFP and mCherry (**fig. 5B**), and infectious virus (**fig. 5D**). In contrast, neurons treated with forskolin at the time of low-dose axonal inoculation produced detectable YFP, but not mCherry (**fig. 5C**), and produced no detectable progeny (**fig. 5D**). These findings suggests that JSH-23 interferes with the establishment of a silent infection in neurons, resulting in lytic replication. However, forskolin treatment is sufficient to induce detectable YFP expression, but infection fails to achieve VP26 expression and lytic replication. These results demonstrate that the dual reporter HSV-1 can achieve latency and reports divergent outcomes of neuronal infection elicited by small molecule manipulation of neuronal signaling.

## Discussion

We sought to observe the temporal progression of HSV-1 replication with single-cell resolution. To achieve this goal, we constructed and isolated a recombinant HSV-1 that would capitalize on the sequential classes of HSV-1 gene expression to report critical transitions in the viral replication cycle. Our dual reporter HSV-1 enables detection of the onset of viral gene expression to the completion of viral replication. In support of the dual reporter HSV-1 retaining wild-type capabilities, we observed comparable replication kinetics, intracellular spread, and competitive fitness in epithelial cell infection (**fig. 1**). We determined that dual reporter HSV-1 demonstrates normal IE and L gene expression that can be observed by YFP and mCherry detection, respectively (**fig. 2**). We employed our model of compartmentalized primary neuron culturing to compare neuronal infection and retrograde-directed spread (**fig. 4**). Comparable extents of replication between unlabeled HSV-1 and dual reporter HSV-1 were observed in neurons. Axonal inoculation of unlabeled HSV-1 or dual reporter HSV-1 resulted in an insignificant difference in the frequency of infected neurons, but a decrease in recovered viral titer from dual-reporter HSV-1-infected neurons was observed. This data suggests that dual reporter HSV-1 retains a capacity for, though a modest reduction in retrograde directed infection. Similarly, there were modest differences in epithelial cell infection between unlabeled and dual reporter HSV-1, particularly regarding plaque size (**fig. 1**). We suspect that propagation and passage of these recombinant viruses has incidentally selected for an enhanced replicative capacity in cultured cells. Parallel analysis of a published single reporter HSV-1 demonstrated similar increase in plaque area compared to unlabeled HSV-1 (**supplemental figure 1**). Despite a potential enhancement in spread of infection, dual reporter HSV-1 exhibited reduced viral fitness in direct competition with unlabeled HSV-1, under conditions of high MOI (**fig. 1**). These experiments provide critical insight into the retained capabilities of the recombinant HSV-1.

The dual reporter HSV-1 visually reports IE and L gene expression to discriminate the progression of viral replication. To evaluate visual outcomes of replication, we assessed fluorescent protein detection in the presence antiviral inhibitors (**fig. 3**). HSV-1-infected cells treated with Acv or PAA exhibited diminished or absent mCherry detection, compared to control treated cells. We observed parallel trends in detection of ICP27 and VP16 by qPCR under these conditions. Interestingly, at 8hpi, detection YFP is not diminished in the presence of Acv or PAA, but ICP27 RNA detection is diminished compared to treated controls. We hypothesize that the incongruence between IE transcript decrease and persistent YFP detection is explained by the CMV promotor regulating YFP expression. While detection of CMV-driven fluorescent cassettes correlate with the onset and extent of IE gene expression (16), the CMV promotor, as a modified beta-herpes virus genetic element may not be responsive to HSV-1 regulation during viral replication. As a result, inhibition of HSV-1 DNA replication did not alter YFP detection at 8hpi, but significantly reduced mCherry-VP26 detection, as well as endogenous IE and L gene expression. Under these conditions, dual reporter HSV-1 reliably reports outcomes of infection that result from aberrant viral replication.

One of the striking applications of the dual reporter HSV-1 was the evaluation of gene expression during the establishment of latent infection in neurons (**fig. 5**). We demonstrate that low-dose axonal inoculation results in no detectable fluorescent reporter detection, consistent with similar models of latent infection (22, 39). Under the same low-dose axonal inoculation conditions, but in the presence of NF-kB inhibitor JSH-23 (40), we observe initial YFP expression and subsequent mCherry detection that steadily increases over a 5-day time course and produces recoverable infectious virus. In contrast, low-dose axonal inoculation in the presence of forskolin, a conventional reactivator of HSV-1 latency *in vitro* (41-44) results in rapid and abundant YFP detection, yet the infected neurons fail to progress to mCherry detection or produce infectious virus. Additionally, spread of infection was not captured over the 5-day course of infection with forskolin treatment, and the number of YFP+ cells diminished. This forskolin-elicited phenotype suggests a ‘stalled’, nonproductive neuronal infection that appears similar to a quiescent model of infection in epithelial cells (15). In this epithelial cell model, CMV-driven FP is robustly detectable, but endogenous IE, E and L HSV-1 transcripts are suppressed (15). Our observations have led to the following hypotheses. First, we hypothesize that low-dose axonal inoculation establishes a latent infection that follows conventional definitions of HSV-1 latency (1, 3, 6, 7). Second, low-dose axonal inoculation in the presence of forskolin yields an intermediate state of infection, allowing transcription from the CMV-driven fluorescent reporter while lytic gene expression from the viral genome remains suppressed, or altered in comparison to lytic replication (45). Further experiments are needed to evaluate the transcriptional profile arising from low-dose neuronal HSV-1 infection with drug treatments. Together, the dual reporter HSV-1 has identified differences in viral gene expression during these altered conditions of neuronal infection.

There are many additional applications for our dual reporter HSV-1 that enable focused investigation of viral replication. Tracking HSV-1 fluorescent reporter detection in epithelial and neuronal cells (**figs. 2 and 5**) provides quantitative measures of viral replication in real time. We can discriminate outcomes of infection based on YFP and mCherry detection. This information is insightful into the progression and outcome of HSV-1 replication at single-cell resolution. Single-cell analyses have highlighted the impressive heterogeneity of HSV-1 infection. HSV-1 gene expression is highly variable within a population of infected cells (28, 47). Additionally, heterogeneity in latent HSV-1 infection of neurons has been investigated (9, 29, 48, 49), and variable states of HSV-1 gnome condensation (50, 51) and gene activity (52) directly influence the capacity for viral reactivation. Expanding temporal observation of dual reporter HSV-1 infected cells using innovative single-cell culturing techniques will provide unmatched insight into HSV-1 gene expression, replication, and pathogenesis (53).

## Materials and methods

### Cell culture

Vero cells purchased from ATCC were maintained in complete DMEM (Dulbecco’s Modified Eagle’s Medium (DMEM) supplemented with 10% fetal bovine serum (v/v) and 1% penicillin/streptomycin). For infection, cells were inoculated with HSV-1 suspended in viral media (DMEM supplemented with 2% fetal bovine serum (v/v) and 1% penicillin/streptomycin. 1 hour-post inoculation, vero cells were aspirated, washed with DPBS, and fed with fresh viral media. When specified, viral media was supplemented with inhibitors (10uM Acyclovir (Fluka PHR1254-1G) or 400ng/mL phosphonoacetic acid (Sigma 4408-78-0) at this timepoint 1 hour-post inoculation. Compounds were prepared and stored at 1000X [stock solution] in DMSO or molecular biology-grade water, respectively.

Mouse Superior Cervical Ganglia (SCG) were excised from embryos at 14 days post gestation from pregnant C57Bl/6 mice. All protocols for isolating SCGs were approved by the Institutional Animal Care and Use Committee (IACUC) at Montana State University (protocol 2022-52-IA). Briefly, isolated SCGs were washed with calcium/magnesium-free HBSS and resuspended in 0.25mg/mL trypsin in calcium/magnesium-free HBSS for dissociation and incubated for 15 minutes in a 37℃ water bath. Trypsinized SCGs were centrifuged and resuspended in 1mg/mL trypsin inhibitor (Gibco 9035-81-8) in calcium/magnesium-free HBSS, then incubated for 5 minutes in a 37°C-water bath. Next, SCGs were centrifuged and resuspended in complete neurobasal and dissociated by trituration using a 5mL Pasteur pipette. After dissociation, neurons were seeded into compartmentalized chambers at approximately 0.7 SCG per chamber. SCG neurons were grown, cultured and infected in complete neurobasal media. 48 hours post seeding, neuronal media was changed and supplemented with 1uM Cytosine-β-D arabinofuranoside (Sigma C6645) for 24 hours. Neuron cultures were fed with fresh complete neurobasal every 3-4 days. Neurons were cultured 21 days to allow maturation and neurite growth prior to HSV-1 infection. For neuronal infection, cells were inoculated with HSV-1 diluted in complete neurobasal media.

### Virus generation and isolation

All viruses used in this manuscript are derived from HSV-1 strain17. All HSV-1 was propagated in vero cells purchased from ATCC. HSV-1 OK13 (HSV-1-YFP) was constructed by homologous recombination of a CMV promoter cassette expressing eYFP with an N-terminal fusion to a mammalian CAAX motif, terminated with SV40 poly(A) signal. The CMV-driven YFP-expression cassette is located between UL37 and UL38 (nt 84,084…84531). HSV-1 OK14 (HSV-1 mCherry-VP26) was constructed by cotransfecting BamHI-digested pHSV1(17+)Lox-mRFPVP26 and purified HSV-1(17+) DNA (O. Kobiler and L. W. Enquist, unpublished data). The pHSV1(17+) Lox-mRFPVP26 was a kind gift from Katinka Döhner and Beate Sodeik. OK15 (dual reporter HSV-1) was isolated by serial isolation of dual-fluorescent plaques arising from HSV-1 OK13 and OK14 coinfection.

### Fluorescence microscopy

Epifluorescence imaging was performed on a Nikon Ti-Eclipse inverted microscope (Nikon Instruments) equipped with a Spectra X LED excitation module (Lumencor) and fast-switching emission filter wheels (Prior Scientific, Rockland MA). Fluorescence imaging used paired excitation/emission filters and dichroic mirrors for yellow fluorescent protein (YFP), GFP and Tetramethylrhodamine (TRITC) channels (Chroma Corp.). All brightfield images were acquired using phase-contrast configuration at 4X, 10X or 20X optical magnification. Vero cells were imaged in 6-well or 12-well plates with conditions performed in triplicate. Primary SCG neurons were imaged in compartmentalized cultures with conditions performed in a minimum of four replicates.

Live imaging was performed on the microscope in conjunction with a stage-top incubation system (Quorum Scientific, Puslinch, Ontario, Canada). Cells were maintained at 37°C in a 5% (vol/vol) CO_2_-enriched atmosphere using a stage-top incubator system. Imaging began 1 hour after application of viral inoculum. Images were acquired every 15 min, and fluorescent illumination intensity was set to 50% power with less than or equal to 100 ms exposure for each channel to avoid photobleaching.

### Time-lapse quantification of YFP and mCherry fluorescence detection

To quantify YFP and mCherry detection, timelapse images were analyzed using Fiji ImageJ (54). First, a circular region of interest (ROI) was drawn around individual cells. The maximum pixel intensity in YFP and mCherry within the determined ROI was then measured for each timeframe. For background subtraction, frame 1 maximum pixel intensity of each ROI was subtracted from the subsequent frames for that ROI. The noise threshold was found to be 32 a.u. for YFP and 120 a.u. for TRITC. N=6 cells presented that were randomly chosen from two fields of view.

### Single-step growth curve of HSV-1 recombinants

Confluent vero monolayers were infected at 10PFU/cell with unlabeled HSV-1, OK13 OK14 or OK15 in triplicate. 1hour-post inoculation, infected vero monolayers were washed with DPBS and fed with viral media. At indicated hpi, cells were harvested via scraping and stored at -80⁰C. Infected cell lysates were assayed for viral titer by plaque assay. Dilutions of infected cell lysates were performed in a 96-well plate and inoculated onto vero cells. 1 hour post-inoculation, vero cells were washed with DPBS and overlayed with viral media supplemented with 1% methylcellulose. Plaque assays were incubated for 72hrs before methylene blue staining for plaque quantification. Each of the triplicate HSV-1 infections was assayed for titer in duplicate.

### Viral competition assay

Confluent vero monolayers were simultaneously infected with unlabeled HSV-1 and OK15 at either 10PFU/cell or 1PFU/cell each virus (21). At 8hpi, infected cells were harvested via cell scraping and stored at -80⁰C. Infected cell lysates were assayed for quantification of viral progeny by plaque assay. Dilutions of infected cell lysates were performed in a 96-well plate and inoculated onto vero cells. 1-hour post-inoculation, vero cells were washed with DPBS and overlayed with viral media supplemented with 1% methylcellulose. At 72hpi, infected cells were imaged, and viral progeny were identified and quantified by plaque fluorescence. Each of the HSV-1 infections were assayed for progeny quantification in triplicate (10PFU/cell n=607, 1PFU/cell n=164).

### HSV-1 plaque area quantification

Confluent vero monolayers in 6 well plates were inoculated with unlabeled HSV-1, OK12 or OK15 at less than 100PFU/well. At 1 hour post-inoculation, cells were washed with DPBS and overlayed with 1% methylcellulose. At 72hpi, infected cells were imaged. Then, infected cells were fixed with cold methanol:acetone and blocked overnight with 2% BSA vol/vol in DPBS. The next day, fixed cells were washed 3x with DPBS then stained with 1⁰ 1:500 Daco antiserum rabbit pAb in 2% BSA vol/vol and incubated overnight. Then, fixed cells were washed 3X with DPBS and given 2⁰ 1:500 donkey anti-rabbit FITC-conjugated mAb overnight. The next day, immunostained cultures were washed 3X with DPBS imaged. Plaque area quantification was performed using NIS Elements analysis polygon measurement tool, measuring the perimeter of plaque-associated fluorescence. Each infection was performed in triplicate, n=200 plaques for unlabeled HSV-1, OK12 and OK15.

### Acyclovir and phosphonoacetic acid

Confluent monolayers of vero cells were infected with OK15 at 10PFU/cell. 1hour-post inoculation, infected vero monolayers were washed with DPBS and fed with viral media that was either supplemented with DMSO, 10uM Acyclovir (Fluka PHR1254-1G) or 400ug/mL phosphonoacetic acid (Sigma 4408-78-0). Compounds were prepared and stored at 1000X stock solution in DMSO or molecular-biology grade water, respectively. Infection was incubated and allowed to proceed for 8 hours before cells were imaged for fluorescent protein detection or harvested for RT-qPCR.

### Quantification of viral gene expression by RT-qPCR

Real-time quantitative PCR of viral transcripts was adapted from (16, 48). Vero cells infected with unlabeled HSV-1 or OK15 at 10PFU/cell and were harvested from 6-well plates at the various timepoints and conditions indicated above using 0.3mL Trizol reagent (Life Technologies), and immediately stored at -20⁰C. Subsequently, RNA was isolated and purified from Trizol following the manufacturers protocol and resuspended in molecular biology-grade water. RNA concentration and quality was assessed by 260/280nm absorbance using an ND1000 UV/Vis NanoDrop (S/N 6612). 2ug RNA was reverse-transcribed using Lunascript RT (NEB E3010L), following the manufacturers protocol. Generated cDNA was stored at -20⁰C. qPCR analysis was performed using PowerUp SYBR Green kit, targeting HSV-1 ICP27 or VP16, or cellular 28S rRNA for endogenous control. Cycle thresholding was auto selected for each target, at approximately Rn = 0.2. Viral transcript abundance is normalized and calculated via ΔΔCt relative quantitation and is represented by log_2_ ^-ΔΔCt^ transformation.

### Single-step growth curve in compartmentalized SCG neurons

Compartmentalized neuronal cultures were inoculated with 10^7^ PFU unlabeled HSV-1 or OK15 on neuronal axons or soma in triplicate. At 3hpi, viral inoculum was removed, and neurons were fed with complete neurobasal. At indicated hpi, neuronal soma were harvested via cell scraping, and stored at -80⁰C. Infected neuronal lysates were assayed for viral titer by plaque assay. Dilutions of infected neuronal lysates were performed in a 96-well plate and inoculated onto vero cells. 1-hour post-inoculation, vero cells were washed with DPBS and overlayed with viral media supplemented with 1% methylcellulose. Plaque assays were incubated for 72hrs before methylene blue staining for plaque quantification. Each of the triplicate neuronal HSV-1 infections was assayed for titer in duplicate.

### HSV-1 VP5 immunostaining and retrograde quantification

Compartmentalized neuronal cultures were inoculated with 10^7^ PFU unlabeled HSV-1 or OK15 on neuronal axons. After 3hpi, viral inoculum was removed, and axons were fed with complete neurobasal media. Infected neuron cultures were incubated, then fixed with cold methanol:acetone at 24hpi. Fixed neuron cultures were blocked overnight with 2% BSA vol/vol in DPBS, washed 3x with DPBS then stained with 1⁰ 1:1000 mouse anti-VP5 mAb in 2% BSA vol/vol overnight. Then, neuron cultures were washed 3X with DPBS and given 2⁰ 1:500 goat-anti-mouse FITC-conjugated mAb overnight. The next day, immunostained cultures were washed 3X with DPBS and counterstained with 1:5000 Hoescht nuclear stain. Cells were washed 1X with DPBS and imaged. DAPI+ and VP5+ neuronal soma were quantified using NIS Elements analysis software. Infections were performed in replicates of 4 neuronal cultures.

